# Genetic and ecomorphological divergence between sympatric *Astyanax* morphs from Central America

**DOI:** 10.1101/2021.04.29.442001

**Authors:** Carlos A. Garita-Alvarado, Marco Garduño-Sánchez, Marta Barluenga, C. Patricia Ornelas-García

## Abstract

Ecological and morphological divergence within populations can be a signal of adaptive divergence, which can maintain intraspecific polymorphisms and promote ecological speciation in the event of reproductive isolation. Here, we evaluate correlations among morphology, trophic ecology, and genetic differentiation between two divergent morphs (elongate and deep-body) of the fish genus *Astyanax* in the San Juan River basin in Central America, to infer the putative evolutionary mechanism shaping this system. We collected the two morphs from three water bodies and analyzed: 1) the correlation between body shape and the shape of the premaxilla, a relevant trophic structure, 2) the trophic level and niche width of each morph, 3) the correspondence between trophic level and body and premaxillary shape, and 4) the genetic differentiation between morphs using mitochondrial and nuclear markers. We found a strong correlation between the body and premaxillary shape of the morphs. The elongate-body morph had a streamlined body, a premaxilla with acuter angles and a narrower ascending process, and a higher trophic level, characteristic of species with predatorial habits. By contrast, the deep-body morph had a higher body depth, a premaxilla with less acute angles, and a broader trophic niche, suggesting generalist habits. Despite the strong correlation between morphological and ecological divergence, the morphs showed limited genetic differentiation, supporting the idea that morphs may be undergoing incipient ecological speciation, although alternative scenarios such as stable polymorphism or plasticity should also be considered. This study provides evidence about the role of ecological factors in the diversification processes in *Astyanax*.

## Introduction

Alternative adaptive phenotypes that utilize different resources within a population (i.e., resource polymorphisms) are commonly found in vertebrates, and they are considered critical in the evolution of innovation and diversification (West Eberhard, 1986, 2003; Smith & Skúlason, 1996; Skúlason et al., 2019). Divergent and frequency-dependent selection can both generate conditions for adaptive divergence by favoring the exploitation of alternative resources, which, in turn, could promote and maintain stable polymorphisms in a population over time (Smith & Skúlason, 1996; West Eberhard, 2003) or, alternatively, promote ecological speciation if divergent phenotypes become reproductively isolated (Schluter, 1996; Schluter, 2000; Nosil, 2012; Hendry, 2009). Genetic processes and/or environmental responses (phenotypic plasticity) can be involved in generating adaptive differences (West Eberhard, 2003; Nonaka et al., 2015).

In the context of ecological speciation, different stages along a continuum can be found (Hendry, 2009; Nosil, Harmon, & Seehausen, 2009), from phenotypic variation without reproductive isolation or stages with a different level of reproductive isolation (incipient speciation) to discontinuous adaptive differences with complete reproductive isolation (speciation completion). It has been argued (e.g., Kulmuni et al., 2020) that the speciation continuum is neither unidirectional nor unidimensional, and that the accumulation of barriers to gene flow is an extended process in which reproductive isolation often slowly evolves. Remarkable cases of resource polymorphisms in trophic ecology and/or habitat use that are considered at different stages of speciation occur in a variety of freshwater fishes. For instance, adaptive divergence without reproductive isolation has been observed in the sunfish *Lepomis gibbosus* (McCairns & Fox, 2004), the yellow perch *Perca flavescens* (Faulks, Svanbäck, Eklöv, & Östman, 2015), the roach *Rutilus rutilus* (Faulks et al., 2015), the brook trout *Salvelinus fontinalis* (McLaughlin & Grant, 1994), and the desert cichlid *Herichthys minckleyi* (Magalhaes, Ornelas-García, Leal-Cardin, Ramírez, & Barluenga, 2015). Intermediate levels of adaptive divergence and reproductive isolation have been reported in sympatric ecotypes of the whitefish *Coregonus clupeaformis* in several North American lakes (Lu & Bernatchez, 1999). Finally, evidence of complete reproductive isolation has been shown in an anadromous species pair of the threespine stickleback (*Gasterosteus aculeatus*) species complex in Japan (Kitano, Seiichi, & Peichel, 2007), and in the Midas cichlid species complex *Amphilophus* spp. in Lake Apoyo, a crater lake in Nicaragua (Barluenga, Stölting, Salzburger, Muschick, & Meyer, 2006).

The complex geological and hydrographical history of Central America makes it a unique scenario for freshwater fish fauna diversification (Myers, 1966; Matamoros, McMahan, Chakrabarty, Albert, & Schaefer, 2015). An excellent system in which to assess mechanisms promoting and maintaining diversity is the freshwater fish genus *Astyanax* Baird & Girard, 1854, given its remarkable adaptability to diverse environmental conditions (Jeffery, 2009). Previous studies have shown the high diversification capacity of *Astyanax* in lacustrine environments (Garita-Alvarado, Barluenga, & Ornelas-García, 2018; Powers et al., 2020), particularly in the San Juan River basin, which includes two large Nicaraguan lakes, Managua and Nicaragua, and their associated outflow river, the San Juan. In this basin, two very distinct morphs (elongate-body and deep-body) of *Astyanax* coexist in sympatry (Garita-Alvarado et al., 2018), raising questions about the degree of morphological divergence, ecological specialization, and reproductive isolation shown by the morphs, and the links between these factors. The divergent morphologies of these two morphs, which have evolved in parallel and coexist in sympatry in other Mesoamerican regions (e.g., Lake Catemaco in Mexico), have been associated with a differential use of environmental resources (Ornelas-García et al., 2018).

Previous morphological studies of *Astyanax* in the San Juan River basin have shown that the alternative morphs present a high level of divergence in trophic-related traits (Garita-Alvarado et al., 2018; Powers et al., 2020). Regarding trophic habits, Bussing (1993), on the basis of a stomach content analysis, described the elongate-body morph as mainly piscivorous, and the deep-body morph as omnivorous, mainly feeding on insects and plant material. However, this analysis represents a single snapshot of trophic habits; therefore, the trophic ecology (trophic niche width and trophic level) of the sympatric morphs in the San Juan River basin is still largely unknown.

Despite the divergence in trophic-related traits (Garita-Alvarado et al., 2018; Powers et al., 2020) and the potential differences in trophic habits (Bussing, 1993), very few studies have characterized the level of genetic differentiation between the two morphs. Ornelas-García, Domínguez-Domínguez, & Doadrio (2008) showed that morphs have a close genetic relationship, and even share mitochondrial haplotypes. Given this context, in the present study, we investigate the degree of genetic differentiation and ecomorphological divergence between the sympatric morphs, and their correlation. We hypothesize that if the two morphs of *Astyanax* are at incipient stages of ecological speciation, then ecomorphological differences would be associated with genetic differentiation, evidencing limitations to gene flow (because of some level of reproductive isolation), as suggested by empirical studies of ecological speciation (Thibert-Plante & Hendry, 2011; Yamasaki et al., 2020). Alternatively, if the two morphs represent a case of stable polymorphisms, we would expect to observe a lack of association between the genetic variation of neutral markers and the ecomorphology of the *Astyanax* morphs due to unrestricted gene flow between them (Smith & Skúlason 1996).

In order to establish whether the morphological, ecological, and genetic divergences between *Astyanax* morphs in the San Juan River basin are associated, we first evaluate the correlation between body shape and trophic morphology (based on the shape of the premaxillary bone shape) to show, in detail, the contrasting patterns of morphological differentiation between the morphs. Then, using stable isotope analyses, we clarify the trophic ecology (trophic niche width and trophic level) of the morphs, and evaluate the association between trophic level and premaxillary and body shape. Finally, by analyzing mitochondrial and neutral nuclear markers, we determine the degree of genetic differentiation between the two divergent morphs.

## Materials and Methods

### Study species and sample collections

We collected the two morphs of *Astyanax* (elongate and deep-body) from ten sampling sites across three water bodies in the San Juan River basin: the two Nicaraguan lakes, Managua and Nicaragua, and the Sarapiquí River (referred to as the three regions throughout the study; Fig. 1). The two morphs were previously considered as species belonging to two different genera; however, phylogenetic studies clearly show that they all belong to the same genus (i.e., *Astyanax*: Ornelas-García et al., 2008; Schmitter-Soto, 2017). Originally, the elongate-body morph was assigned to the genus *Bramocharax*, and comprised two species: *B. bransfordi* Gill 1877 from Lake Nicaragua and the Sarapiquí River, and *B. elongatus* Meek 1907 from Lake Managua. The deep-body morph was assigned to the genus *Astyanax*, and also comprised two species: *A. nicaraguensis* Eigenmann & Ogle 1907 from Lake Nicaragua and the Sarapiquí River, and *A. nasutus* Meek 1907 from Lake Managua. We collected a total of 302 specimens: 187 deep-body morphs and 115 elongate-body morphs. The number of specimens and sampling sites included varied depending on the analysis (see below). The low sample size of the elongate-body morph from Sarapiquí River reflects its low abundance in lotic habitats, where its presence is considered atypical (Bussing, 1998). Fish were collected using either gill or cast nets during sampling campaigns that took place between November 2011 and January 2017. The subsample of fish used in the isotopic analyses were collected during the dry season between December 2016 and January 2017. While in the field, the collected fish were euthanized with iced water, photographed in a standardized position, and assigned to a morph type on the basis of dentition patterns (Bussing 1998). Fin clips and muscle tissue samples were collected and preserved for, respectively, the DNA analyses and the stable isotope analyses. The muscle samples were stored in sodium chloride powder in the field, and later at -80 °C in the laboratory. Finally, the premaxillary bone was dissected for the morphometric analyses.

**Figure 1.**
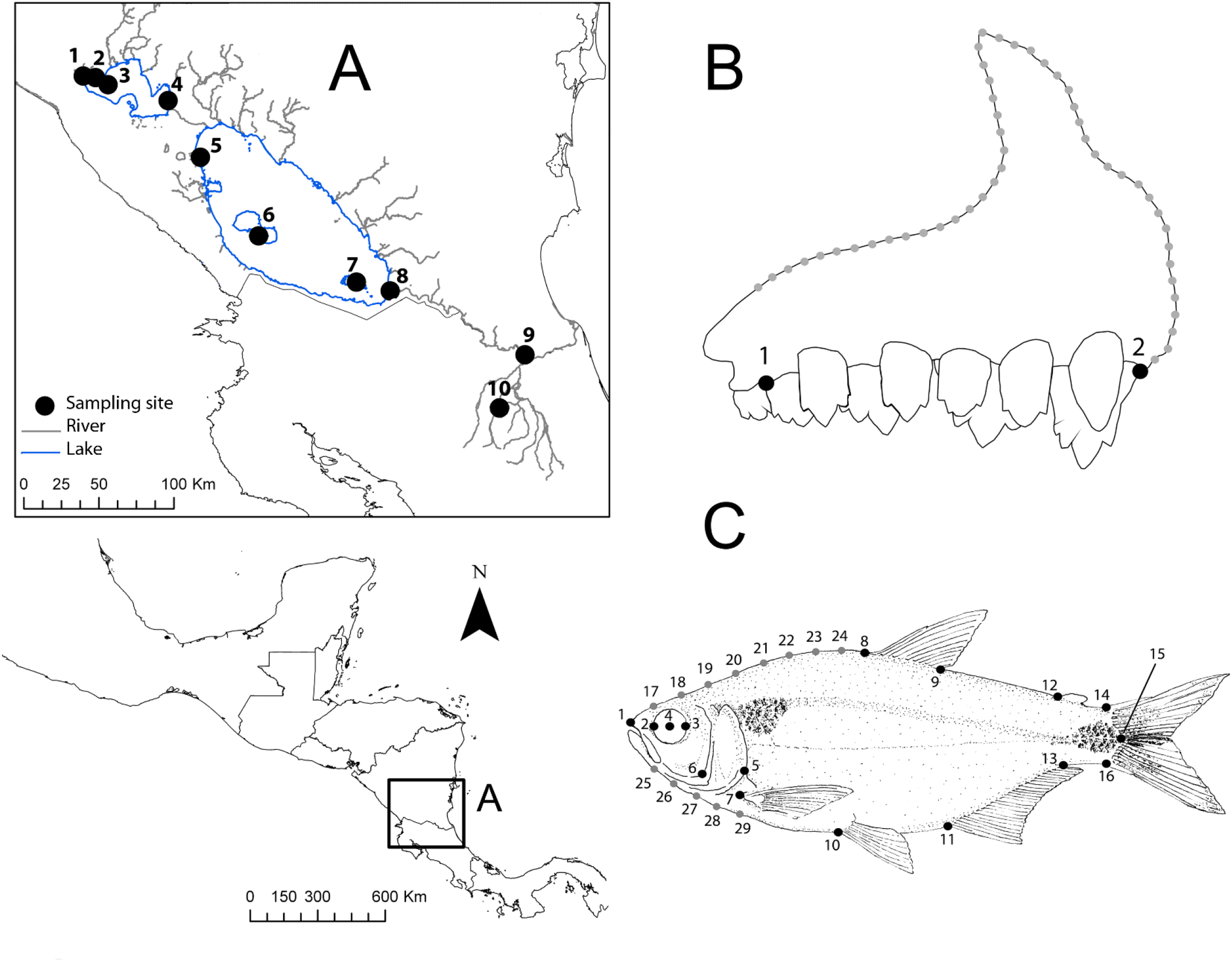
A) Map representing sampling sites in the San Juan River basin in Central America. Sampling localities 1–4, Lake Managua; 5–8, Lake Nicaragua; and 9 and 10, Sarapiquí River. B) Distribution of landmark (black) and semilandmark (grey) positions used to characterize the premaxilla bone. Landmark positions: 1) the posterior insertion of the penultimate tooth, and 2) anterior insertion of the anterior tooth. Semilandmark positions: 50 equally distributed positions describe the dorsal profile of the premaxilla from the anterior tooth to the projection of landmark 1 in the dorsal profile of the premaxilla bone. C) Distribution of landmark (black) and semilandmark (grey) positions to characterize the body shape. Landmark positions: 1) snout tip, 2) anterior extent of eye, 3) posterior extent of eye, 4) center of eye, 5) posterior extent of the operculum, 6) lower margin of preoperculum, 7) anterior insertion of pectoral fin, 8) anterior and 9) posterior insertion of dorsal fin, 10) anterior insertion of pelvic, 11) anal, and 12) adipose fin, 13) posterior insertion of anal fin, 14) dorsal insertion of caudal fin, 15) posterior extent of caudal peduncle, and 16) ventral extent of caudal fin and caudal peduncle. Semilandmark positions: eight equally distributed positi ons between dorsal projection in body profile of the anterior extent of eye and the same projection of the pelvic fin (semilandmarks 17 to 24), and five equally distributed positions between ventral projection in body profile of the anterior extent of eye and the same projection of the anterior insertion of the pectoral fin (semilandmarks 25 to 29).

### Ecomorphology

To characterize the shape of the premaxillary bone and the body, we performed geometric morphometric analyses. Prior to imaging, the dissected premaxillary bone was cleaned in a 1 M KOH solution for 15 min. We imaged each premaxilla using a stereoscope (Zeiss Stemi305), and digitized two landmarks and 50 semilandmarks (Fig. 1B) using TPSDig2 version 2.31 (Rohlf, 2015). We performed a Procrustes superimposition, and calculated relative warps (RWs) and centroid size using the program TPSRelw version 1.69 (Rohlf, 2015). To characterize body shape, we digitized 16 landmarks and 13 semilandmarks (Fig. 1C) also using TPSDig2. We performed a Procrustes superimposition, and calculated RWs and centroid size with CoordGen8h and PCAGen8o, respectively, in the Integrated Morphometrics Package (Sheets, 2001).

To test for an association between trophic morphology and body shape, and to characterize in greater detail the morphological divergence between morphs, we correlated the shape of the premaxilla with body shape for each morph through an analysis of 136 specimens (92 deep-body and 44 elongate-body morphs from eight of the 10 sites). We used a linear mixed-effects model (LMM) to evaluate the association of RW1 of the premaxilla, which is directly related to ecomorphological trophic specialization (as the response variable), with morph type, RW1 of body shape, and region (as fixed effect predictor variables; only statistically significant interactions were included). We included sampling site as a random effect. The LMM was run in the nmle package in R (R Core Team, 2013, version 3.03: http://cran.r-project.org).

### Trophic divergence analysis

To assess trophic divergence between morphs, we analyzed the stable isotope ratios of carbon (13C/12C) and nitrogen (15N/14N) in 96 samples (59 deep-body and 37 elongate-body) collected from seven of the 10 sampling sites. These ratios are widely used to determine trophic level and trophic niche width (Post, 2002; Jempsen & Winemiller, 2002; Layman, Arrington, Montaña, & Post, 2007; Schalk, Montañ a, Winemiller, & Fitzgerald, 2017). Preserved muscle tissue samples were dried at 60 °C for 24 h in an oven and then ground into a fine powder with a mortar and a pestle. Subsamples (1–1.5 mg) were individually packaged into tin capsules prior to being analyzed at the Stable Isotope Facility of the University of California, Davis. The stable isotopic ratios were measured in parts per thousand (‰) in standard delta “δ” notation. To determine trophic niche width, we used metrics that describe trophic variation based on the position of consumers in δ13C and δ15N (Layman et al., 2007; Jackson, Inger, Parnell, & Bearhop, 2011). For each morph in each of the three regions, we calculated six trophic niche metrics (δ15N range, δ13C range, total area: TA, mean distance to centroid: CD, mean nearest neighbor distance: NND, and standard deviation of nearest neighbor distance: SDNND) following Layman et al. (2007). We also calculated the Bayesian standard ellipse area (SEA_B_) using the package SIBER (Jackson et al., 2011) in R. Similar to TA (Layman et al., 2007), SEA_B_ provides an estimate of the trophic niche width per morph, however, it is less influenced by sample size compared with TA (Jackson et al., 2011; Syvä ranta, Lensu, Marjomä ki, Oksanen, & Jones, 2013).

We next compared the trophic level of morphs by region, and its relationship to the shape of both the premaxilla and the body. For this, we analyzed the shape of the premaxilla in 70 specimens, and the shape of the body in 96 specimens. First, we visualized variation between morphs within regions in these shapes by plotting RW2 against RW1. Then, we performed two LMM analysis using δ15N as the response variable and either RW1 of the premaxillary shape or of the body shape as covariate. In each analysis, we included morph type, the covariate, and region as fixed effect predictor variables (we tested for an interaction between all predictor variables but included only if statistically significant). We included sampling site as a random effect.

Size influences many traits in animals including shape (Gould, 1966; Klingenberg, 1996). To control for size and to determine the variation in the shape of both the premaxilla and body associated with size (centroid size), we performed independent LMM analyses using the RW1 of premaxillary shape or of body shape as the response variables. The log centroid size, morph type and region were the fixed effect predictors. For both analyses, sampling site was included as a random effect. All of the LMM analyses were performed using the package nlme in R.

### Genetic analyses

DNA was extracted using a standard Proteinase-K in a SDS/EDTA solution, NaCl and chloroform, following the protocol by Sonnenberg, Nolte, and Tautz (2007). DNA quality and quantity were determined with a Nanodrop 1000 (Thermo Fisher Scientific). A fragment of 880-bp of the mitochondrial gene cytochrome *b* (Cyt*b*) was amplified by PCR from 37 specimens from eight sampling sites using the primers Glu- F 5’GAAGAACCACCGTTGTTATTCAA and Thr-R 5’ACCTCCRATCTYCGGATTACA (Zardoya & Doadrio, 1998). The following thermocycling conditions were used: initial denaturation at 95 °C (5 min), followed by 35 cycles at 94 °C (45 s), 48 °C (1 min), and 72 °C (90 s), and a final extension at 72 °C (5 min). Each PCR was performed in a final volume of 10 µl containing 0.4 mmol of each primer, 0.2 mmol of each dNTP, 2 mmol MgCl2, 1 unit of Taq DNA polymerase (Invitrogen), and 10 ng of template DNA. PCR products were purified with EXOSAP-IT (Thermo Fisher Scientific) and then sequenced on an ABI PRISM 3700 DNA automated sequencer (Applied Biosystems).

We carried out a phylogenetic reconstruction of *Astyanax* spp. in Central America (i.e., Group II in Ornelas-García et al., 2008), we used 37 sequences generated in this study and 59 that were previously published by Ornelas-García et al. (2008) using an alignment of 1140-bp. The latter sequences represent populations of *Astyanax* from tributaries near the Pacific coast of northern Nicaragua, the Atlantic coast of Nicaragua and Costa Rica and northern Central America. We included lineages of *Astyanax* from Mexico and Costa Rica (Groups I and III, respectively) as the outgroups. The Bayesian inference analysis was performed in MrBayes version 3.2.7 (Huelsenbeck & Ronquist, 2001), as implemented in the CIPRES Science Gateway (https://www.phylo.org/portal2/home.action; Miller, Pfeiffer, & Schwartz, 2010). We created codon partitions for the first, second, and third position in the alignment, and used GTR + Γ as the substitution model. The phylogenetic reconstruction was performed using two independent runs of four Metropolis-coupled Markov chain Monte-Carlo for 20 million generations each, with the first 20% of trees discarded as burn in. The sampled trees were used to estimate posterior probabilit ies and determine branch supports. We assessed convergence of the Bayesian analysis in MrBayes, and also confirmed that ESS values were well over 200 using Tracer (Rambaut, Drummond, Xie, Baele, & Suchard, 2018).

We amplified 11 nuclear microsatellite loci that had been developed for *Astyanax mexicanus* (Protas, Conrad, Gross, Tabin, & Borowsky, 2006) from 302 specimens (187 deep-body morphs, 115 elongate-body morphs) across all sampling sites. These markers are located on different chromosomes and are not linked to reported traits. The loci were multiplexed in two reactions, each with a final volume of 5 μl, using the Multiplex PCR Kit (QIAGEN), following the manufacturer’s instructions. PCR amplifications consisted of initial denaturation at 95 °C for 5 min; followed by 35 cycles of denaturation at 94 °C for 30 s, annealing at 52–60 °C for 30 s, and extension at 72 °C for 45 s; and a final extension step at 72 °C for 7 min. Forward primers were labelled with fluorescent dyes (Invitrogen), and amplified PCR products were run on an ABI Prism 3730 DNA Analyzer (GS500 ROX size standard). Allele sizes were scored using Geneious R10 (Kearse et al., 2012), and MICRO-CHECKER version 2.23 (Van Oosterhout, Hutchinson, Wills, & Shipley, 2004) was used to check for the presence of null alleles and to identify and correct any genotyping errors.

For the genetic differentiation analyses and summary statistics for Cyt*b* we included an alignment of 880-bp and a total of 49 sequences: 37 generated in this study and 12 previously published ones of *Astyanax* from the San Juan River basin (Ornelas-García et al., 2008). The microsatellite analysis included 302 samples of *Astyanax*. Summary descriptive statistics were calculated for combinations of morphs and regions using DnaSP version 6 (Rozas et al., 2017) for Cyt*b* and GENALEX version 6 (Peakall & Smouse, 2006) for the microsatellites. For the microsatellites, allelic richness (AR), using the rarefaction approach to deal with differences in sample size, was calculated in the package popgenreport (Adamack & Gruber, 2014) in R. We evaluated the degree of genetic differentiation between morphs and between geographical regions using several methods. Initially, for both types of markers, we performed two hierarchical analyses of molecular variance (AMOVAs) each with 20,000 permutations in Arlequin version 3.5.2.2 (Excoffier & Lischer, 2010). The first included morphs as groups (and regions as populations), and the second considered regions as groups (and morphs as populations). Then, we calculated the pairwise Fst of the six morph type–region combinations (for both types of molecular markers), also in Arlequin, with 20,000 permutations.

To determine the genetic clustering of morphs or regions using nuclear data, we employed two methods. First, we ran a Bayesian model-based clustering algorithm in Structure version 2.1 (Pritchard, Stephens, & Donnelly, 2000) to test the assignment of individuals according to morph type and region. The admixture model and the option of correlated allele frequencies between populations were used. To determine the number of ancestral clusters, K, we compared the log-likelihood ratios of five runs for values of K between 1 and 11, and calculated the ΔK following Evanno, Regnaut, and Goudet (2005) to determine the K value associated with the best log-likelihood probability. Each run consisted of 1,000,000 iterations with a burn-in period of 200,000 without prior information on their sampling region. Also, we ran the clustering algorithm of Structure to test for any particular genetic structuring for each region separately, including the same parameters used in the overall Structure analysis for values of K between 1 and 6. Second, we performed a discriminant analysis of principal components (DACP) (Jombart & Ahmed, 2011; Jombart & Collins, 2015) using the package adegenet in R. The DA method defines a model in which genetic variation is partitioned into a between-group and a within-group component, yielding synthetic variables that maximize the first group while minimizing the second (Jombart & Ahmed, 2011). The DAPC was performed using the microsatellite data, with missing data removed, and the individual clustering assignment was determined using the *find*.*clusters* function and a sequential K-means from 1 to 6 with 10 iterations. The best-supported model inferring the optimal number of genetic clusters was selected on basis of the Bayesian Information Criterion (BIC). The validation set was selected with a stratified random sampling, which guarantees that at least one member of each conglomerate or cluster is represented in both the training and the validation set (Jombart & Collins, 2015). The proportions of intermixes, obtained from the membership probability based on the retained discriminant functions, were plotted for each individual.

Finally, to test for isolation by distance, we pooled the specimens by sampling site and performed a Mantel test on the microsatellite data in Arlequin with 20,000 permutations.

## Results

### Ecomorphology

The LMM analysis testing for an association between trophic morphology and body shape revealed a strong correlation between the first RWs of both premaxillary and body shape (RW1 of body, F_1,124_ = 19.3, p < 0.001). We also observed clear morphological differences between the morphs (F_1,124_ = 420.3, p < 0.001). Positive RW1 values for body shape, indicating slender bodies, were associated with positive RW1 values for premaxillary shape, which depict premaxillas with acute angles and a narrow ascending process, both characteristic of the elongate-body morph (Fig. 2). Despite some overlap in body shape between morphs (between 0 and 0.02 on the x-axis, Fig. 2), its relationship with the shape of the premaxilla clearly differed between morphs. We also found marginal differences among regions (F_2,5_ = 5.8, p = 0.05), as well as different slopes (interaction region*RW1 of body, F_2,124_ = 4.7, p < 0.05).

**Figure 2.**
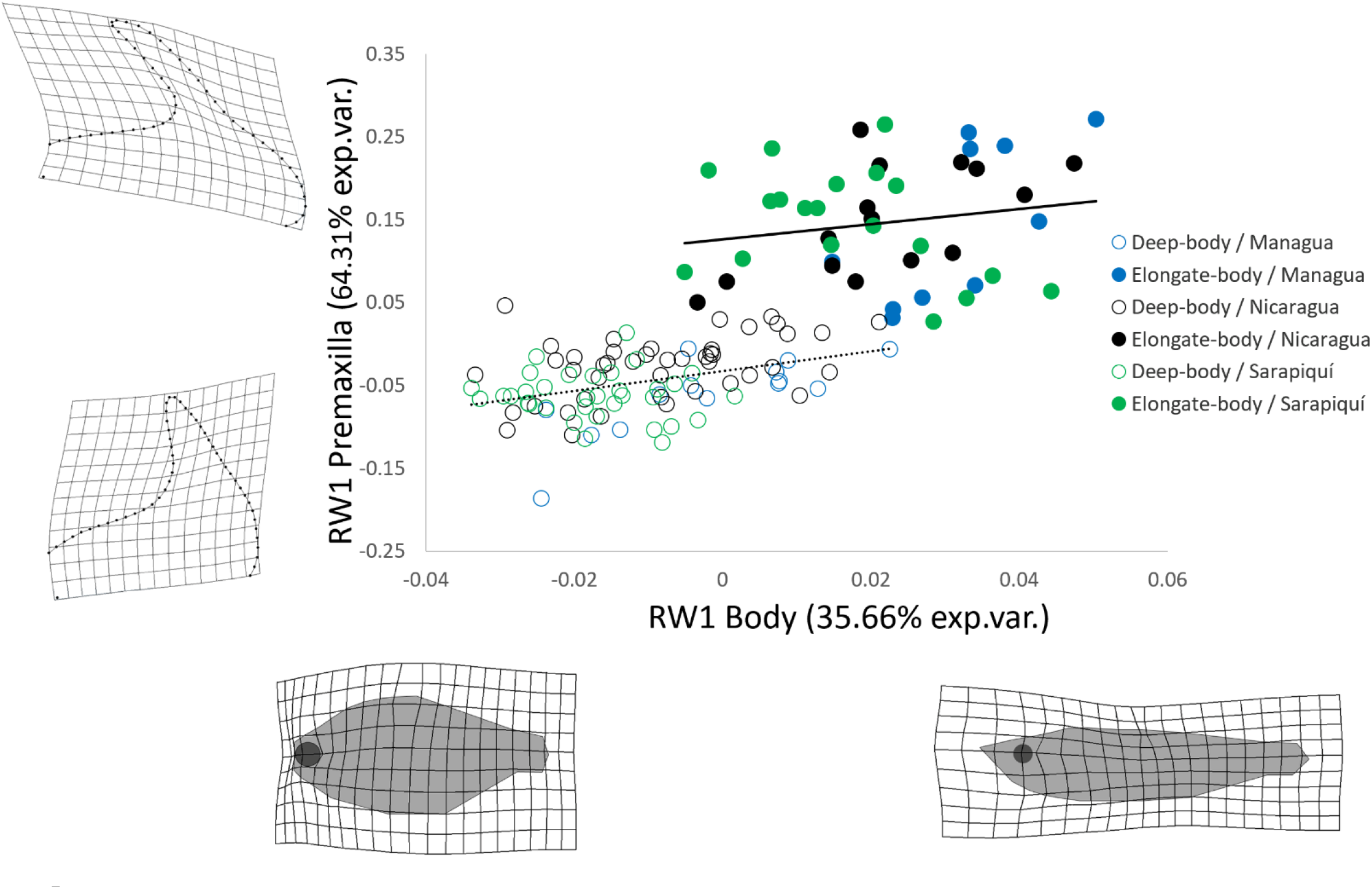
Relationship between RW1 of premaxillary shape and RW1 of body shape by region and morph type. The solid line indicates the elongate-body morph, and the dotted line, the deep-body morph. The empty circles correspond to the deep-body morph, and the filled circles, the elongate-body morph. Thin-plate spline transformations of the landmark positions represent the extreme transformation of the premaxilla shape along the y-axes, and the body shape along the x-axes.

### Trophic divergence

The results of the stable isotope ratio analysis showed that sympatric morphs within regions differed in trophic niche width, although the direction of divergence was region dependent (Fig. S1A). In the lakes Managua and Nicaragua, the trophic niche width of the deep-body morph was broader than that of the elongate-body morph according to the isotope metrics (Fig. S1A); moreover, the TA, CD, and SEAB metrics were greater for the deep-body morph (Fig. S1B, Table S1). In contrast, in the Sarapiquí River, the elongate-body morph showed a greater trophic niche width compared with the deep-body morph, mainly due to having a larger δ13C range (Fig. S1A, Table S1); however, our sample size was low for the calculation of this trophic niche.

When RW2 of the premaxillary shape was plotted against RW1, sympatric morphs could be clearly discriminated for all regions (Fig. 3A). RW1 mainly accounted for the variation in the angles and the width of the ascending process. Acute angles and a narrow ascending process are associated with the elongate-body morph and a higher trophic level (with the deep-body morph showing the opposite patterns). RW2 accounted for the variation in the width of the premaxilla and the length of the ascending process. For body shape, RW1 mainly discriminated individuals by body depth, while RW2 was related to head length and eye diameter (Fig. 3B). Sympatric morphs from Lake Managua and Sarapiquí River were clearly distinguishable by body shape, whereas those from Lake Nicaragua showed some overlap, but followed the same pattern observed for the other two regions.

**Figure 3.**
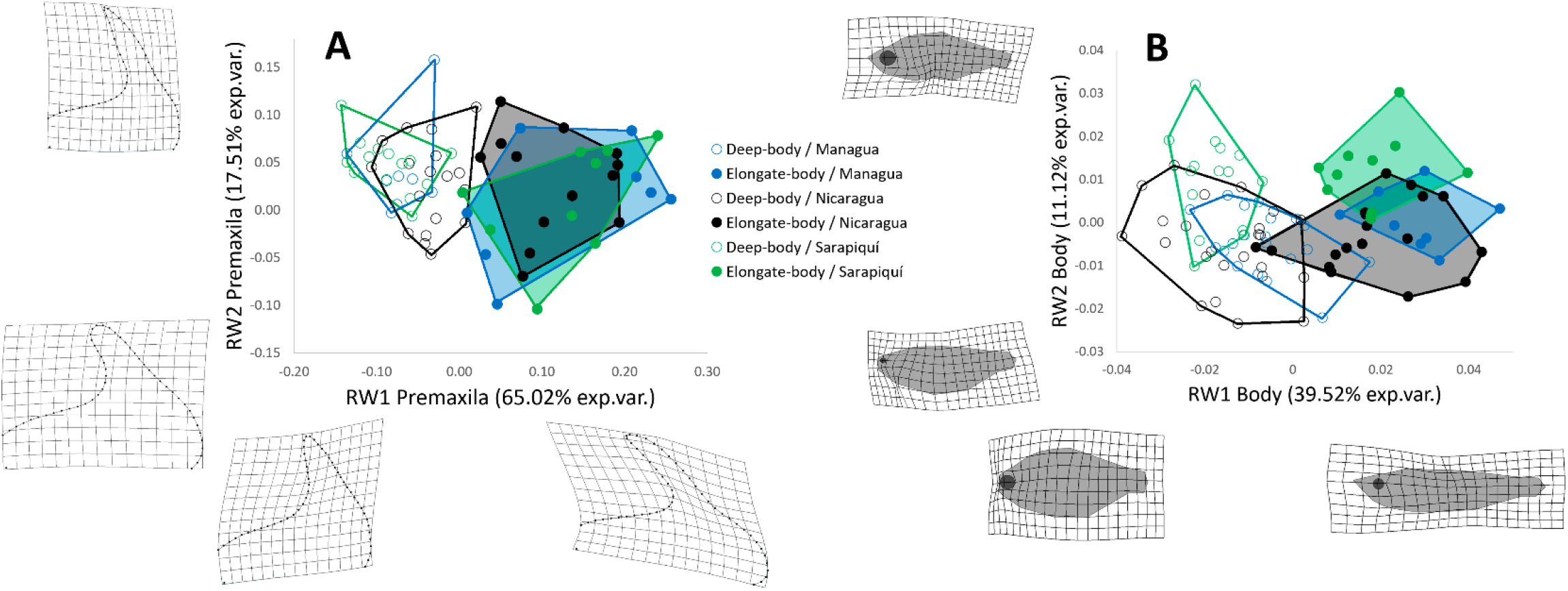
Relative warp analysis of the shape of the premaxilla (A) and body shape (B). Thin-plate spline transformation of the landmark positions represent the e xtreme transformation along the axes. The empty circles correspond to the deep-body morph, and the filled circles correspond to the elongate-body morph.

In the comparisons of trophic levels between morphs, the elongate-body morph showed a higher trophic level than the deep-body morph, when both premaxillary shape and body shape were considered (Fig. 4). In the LMM with the RW1 of premaxillary shape set as the covariate, we found significant differences between morph types (F_1,61_ = 32.9, p < 0.001) and regions (F_2,4_ = 74.2, p < 0.01), and the RW1 of the premaxillary shape was positively correlated with δ15N values (Fig. 4A–C, F_1,61_ = 5.3, p < 0.05). Likewise, in the LMM with the RW1 of the body shape set as covariate, we found significant differences associated with morph type (with the elongate-body morph showing a higher trophic level, F_1,85_ = 25.3, p < 0.001) and regions (F_2,4_ = 24.2, p < 0.01), and a positive correlation between the RW1 of the body shape and δ15N (Fig. 4D–F, F_1,85_ = 4.8 p < 0.05). The extent to which the two morph types differed was region dependent, with morphs from Lake Nicaragua showing a smaller divergence (interaction region*morph F _2,85_ = 8.9, p < 0.001) than those from Lake Managua and the Sarapiquí River. Regarding shape variations associated with size, differences between the two morph types in the RW1 of both the premaxillary shape and the body shape were consistent and independent of variations in size (log centroid size) (Fig. S2).

**Figure 4.**
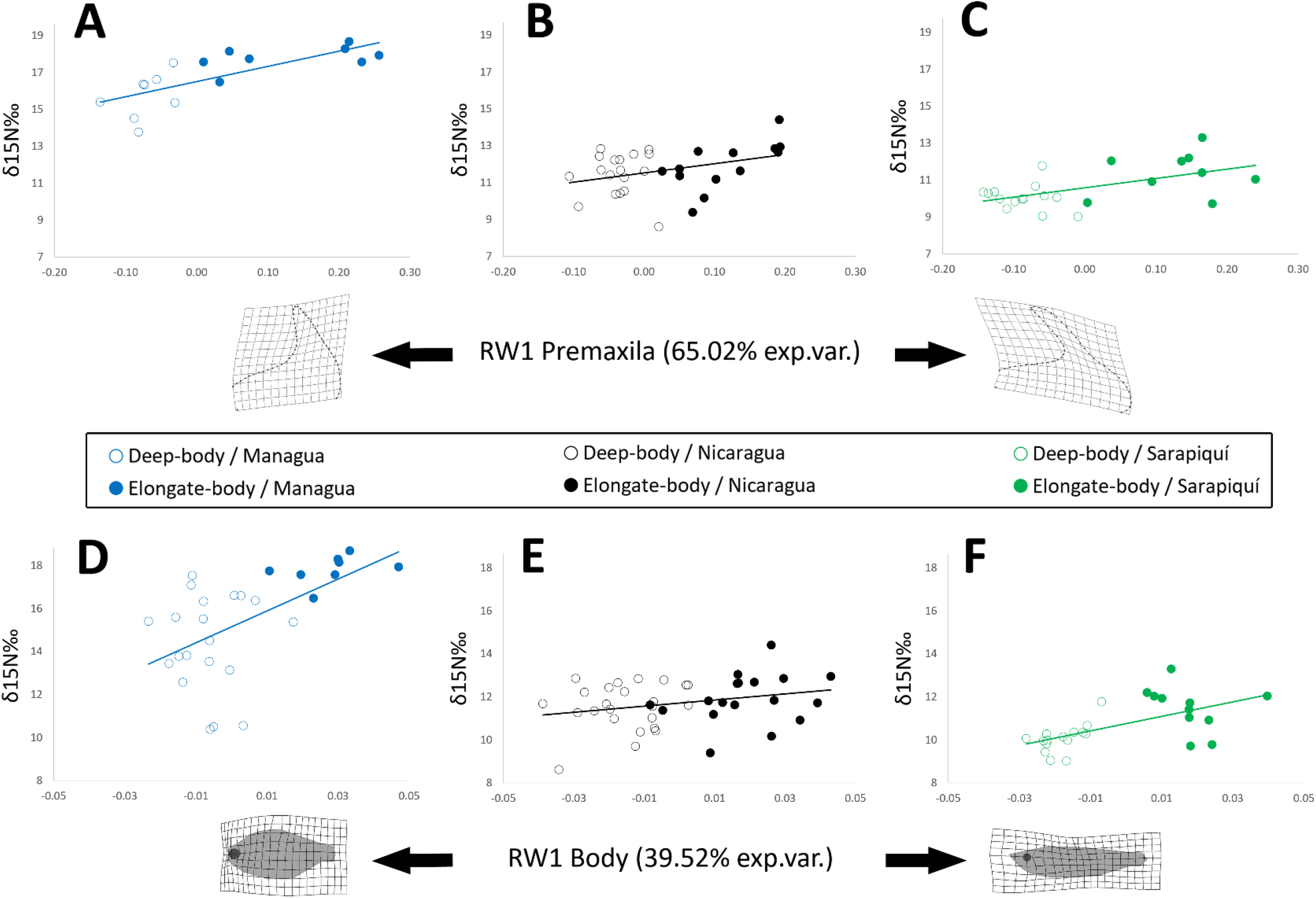
Relationship between trophic level (δ15N) and the shape of the premaxilla (A–C) and the body (D–F), measured by RW1 of the morphs in the three main regions.

### Genetic analyses

We recovered five well supported clades in the Bayesian phylogenetic tree based on Cyt*b* of Central American *Astyanax* (Fig. S3). The relationships among these clades were not fully resolved, and the morphs were not recovered as a monophyletic group. Two of the clades (A and B) comprised samples from the three regions in the San Juan River basin. Clade A included samples of both morphs from Lake Managua and those from tributaries along the Pacific coast of northern Nicaragua. Clade B was comprised of samples of both morphs from all three regions in the San Juan River basin as well as those from tributaries along the Atlantic coast of Nicaragua and Costa Rica. The other clades were comprised of samples from northern Central America (Clades C and D) and the Pacific versant of Costa Rica (Clade E).

Regarding the nuclear markers, of the 11 microsatellite loci analyzed, some showed signals of the presence of null alleles in some of the studied populations, and only one (16C) showed an excess of homozygotes in all groups considered. Summary statistics of the genetic variability of Cyt*b* and the microsatellites are shown in Table 1. For Cyt*b*, we found high levels of polymorphisms in the six groups analyzed (i.e., the two morph types in each of the three regions). These groups also showed high levels of haplotype diversity (HD, 0.9–1), and a high proportion of private haplotypes (PH) (40 %–85.7%; Table 1A). All microsatellites were polymorphic in all groups. The deep-body morph from Lake Managua had the most alleles (Na) and effective alleles (Ne), while the elongate-body morph from Sarapiquí River had the fewest alleles. However, after controlling for sample size, the deep-body morph from Sarapiquí showed the highest allelic richness (Table 1B). Also, observed (Ho) and expected heterozygosity (He) and unbiased expected heterozygosity (UHe) were relatively high for all groups. The elongate-body morph from Sarapiquí River exhibited the highest value of Ho, and the deep-body morph from the same river showed the highest value of He (Table 1B).

**Table 1.** Summary statistics for the genetic variability of mitochondrial (Cyt*b*) (A) and 11 nuclear microsatellite loci (B) for the two morphs of *Astyanax* from the three studied regions of the San Juan River basin. n = sample size, h = number of haplotypes, Hd = haplotype diversity; Pi = nucleotide diversity; PH = % of private haplotypes; Na = number of alleles; Ne = number of effective alleles, AR = allelic richness, Ho = observed heterozygosity; He = expected heterozygosity; UHe = unbiased expected heterozygosity.

|  | L. Nicaragua |  | L. Managua |  | R. Sarapiquí |  | All |
| --- | --- | --- | --- | --- | --- | --- | --- |
|  | Deep | Elongate | Deep | Elongate | Deep | Elongate |  |
| A) *Cytb* |  |  |  |  |  |  |  |
| n | 14 | 6 | 13 | 4 | 7 | 5 | 49 |
| h | 11 | 5 | 10 | 4 | 7 | 4 | 33 |
| Hd | 0.95 | 0.93 | 0.95 | 1 | 1 | 0.90 | 0.96 |
| Pi | 0.01 | 0.01 | 0.02 | 0.02 | 0.01 | 0.01 | 0.02 |
| PH | 72.7 | 40 | 80 | 50 | 85.7 | 75 | - |
| B) Microsatellites |  |  |  |  |  |  |  |
| n | 72 | 68 | 82 | 35 | 33 | 12 | 302 |
| % polymorphic loci | 100 | 100 | 100 | 100 | 100 | 100 | 100 |
| Na | 18.09 | 17.45 | 20.27 | 15.00 | 15.91 | 8.82 | 15.92 |
| Ne | 8.78 | 9.01 | 10.69 | 8.78 | 10.52 | 5.71 | 8.91 |
| AR | 9.99 | 9.67 | 10.81 | 9.82 | 10.86 | 7.93 | - |
| Ho | 0.65 | 0.64 | 0.68 | 0.67 | 0.69 | 0.70 | 0.67 |
| He | 0.85 | 0.83 | 0.87 | 0.84 | 0.88 | 0.80 | 0.84 |
| UHe | 0.86 | 0.84 | 0.87 | 0.85 | 0.90 | 0.83 | 0.86 |

In the hierarchical AMOVA s for Cyt*b* and the microsatellites, similar results were obtained when samples were grouped by morphs, with significant differences found among regions (i.e., Lake Managua, Lake Nicaragua, and Sarapiquí River). Also, in the AMOVA for Cyt*b*, when samples were grouped by region, only the within populations level was significant (Table 2A). However, in the AMOVA for the microsatellites, we found significant differences between morphs when samples were grouped by region (Table 2B). In all groupings, the within population level was significant, and accounted for most of the explained variance, as frequently seen in AMOVA (Excoffier, Smouse, & Quattro, 1992) (Table 2).

**Table 2.** Two-level AMOVA for Cyt *b* (A) and the microsatellites (B). Bold values indicate significant differences (p < 0.05).

|  | Among groups |  | Among populations within groups |  | Within populations |  |
| --- | --- | --- | --- | --- | --- | --- |
|  | Fct | % variation | Fsc | % variation | Fst | % variation |
| **A) *Cytb*** |  |  |  |  |  |  |
| **Regions** (Managua, Nicaragua, and Sarapiquí) | 0.13 | 13.91 | -0.016 | -1.39 | **0.12** | 87.48 |
| **Morphs** (elongate and deep-body) | -0.013 | -1.34 | **0.109** | 11.06 | **0.097** | 90.28 |
| **B) microsatellites** |  |  |  |  |  |  |
| **Regions** (Managua, Nicaragua, and Sarapiquí) | 0.006 | 0.63 | **0.008** | 0.85 | **0.014** | 98.52 |
| **Morphs** (elongate and deep-body) | 0.0006 | 0.06 | **0.013** | 1.3 | **0.013** | 98.64 |

In the Fst pairwise comparisons of the mitochondrial data, no significant differences were found (Table 3). However, for the nuclear data, significant differences between sympatric morphs were observed only for the Sarapiquí River. Significant differences between other pairwise comparisons of morphs and regions are shown in Table 3.

**Table 3.**
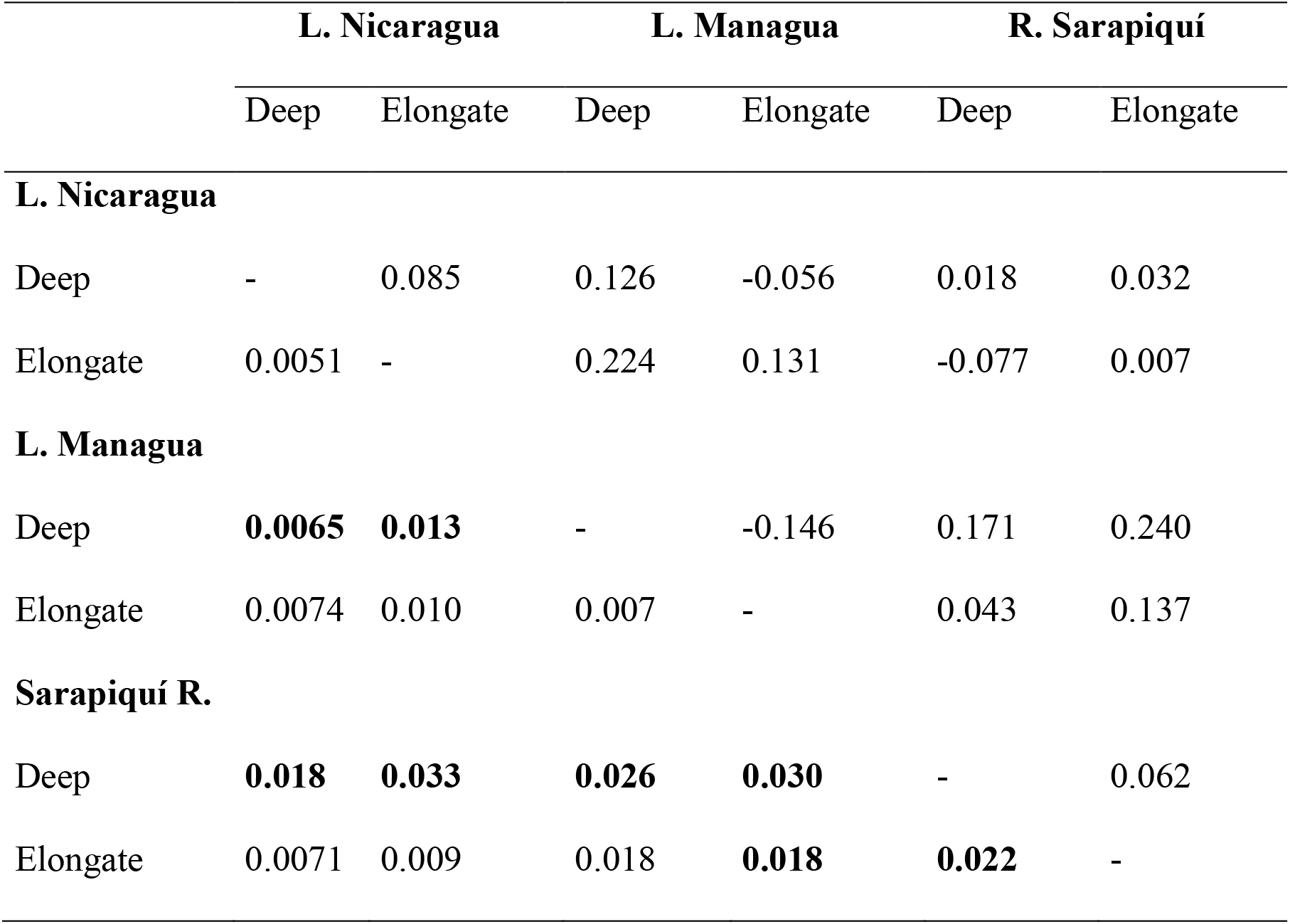
Pairwise Fst comparisons across morphs and regions. Above the diagonal, Cyt*b* and below the diagonal, microsatellite data. Bold values indicate significant differences after Bonferroni correction.

In the Structure analysis and according to the ΔK value following Evanno et al. (2005), a K of 2 showed the best log-likelihood probability. However, the clusters did not group according to morph type or region (Fig. S4A). Clusters also did not group by morph type in the analysis by region (Fig. S5). In the DAPC, we retained the first 60 principal components, and the BIC indicated the best-fit k value was 3; however, as in the Structure analyses, clusters did not group according to morph type or region (Fig. S4B).

The Mantel test showed an isolation by distance pattern, evidencing that genetic distances between sampling sites among regions were larger than those between sampling sites within regions (r^2^ = 0.45, p < 0.001, Fig. 5).

**Figure 5.**
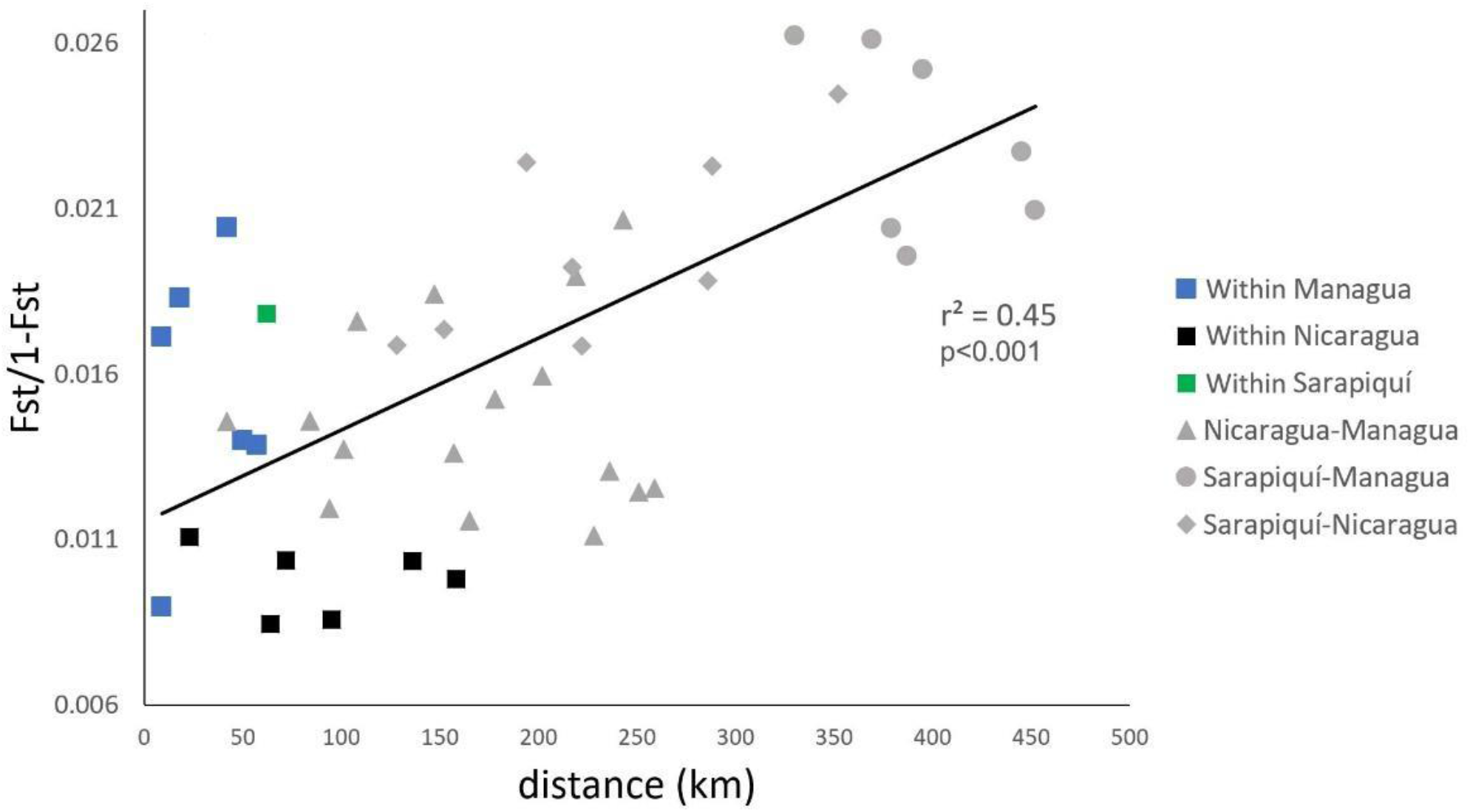
Isolation by distance (IBD) pattern in San Juan River system as shown by the Mantel test.

## Discussion

In this study, we evaluated the association among morphological, ecological, and genetic divergences between *Astyanax* morphs across three tropical regions of the San Juan River basin in Central America. Our results showed an association between ecological divergence in trophic niche and trophic level (i.e., resource use) and morphological differentiation in the body and premaxillary shape in *Astyanax* (trophic polymorphism), which could be interpreted as evidence of adaptive divergence. Moreover, we observed a weak but significant level of genetic differentiation in the nuclear markers between the sympatric morphs (AMOVA and pairwise Fst in Sarapiquí River). Although we cannot rule out a possible scenario of stable trophic polymorphism, the evidence of slight genetic differentiation provides some support to the hypothesis that morphs may be in an incipient speciation process. Regardless, our study provides strong additional evidence about the role of ecological factors in the diversification processes in *Astyanax*.

### Morphological and ecological divergence between morphs

Sympatric morphs of *Astyanax* from the San Juan River basin showed a high level of morphological divergence when correlating body shape with premaxillary shape. As body shape becomes more streamlined, the angles of the premaxilla become acuter and the ascending process narrower (note the almost parallel lines in Fig. 2). However, it is the difference in the magnitude of the premaxillary shape (i.e., the separation of morph-adjusted lines along the y-axes) that clearly separate the two morphs, despite some overlap in body shape. Modularity, i.e., the presence of discrete modules as sets of phenotypic traits that are tightly integrated internally but relatively independent from other modules (Klingenberg, 2008, 2014), has been recognized as a universal emergent property of biological traits that could facilitate evolution (West-Eberhard, 2019). Indeed, modularity has already been suggested to be involved in the divergence of lacustrine *Astyanax* in other lineages (Ornelas-García, Bautista, Herder, & Doadrio, 2017). Notably, in our case, the results demonstrate a strong correlation between the shape of the premaxilla and body shape in morphs of Central American *Astyanax*, revealing contrasting patterns of covariation between them. Therefore, future analyses of the modular evolution of both morphs may shed light on the synergistic role of the two morphological sets (body shape and the trophic morphology) in the divergence of the morphs, and the role of modular evolution as a triggering mechanism in the trophic diversification of the genus.

The strong correlation between body shape and the shape of the premaxilla in these morphs highlights the relevance of ecomorphology, along with differences in trophic ecology, as a promoter of diversification in *Astyanax*. For example, slender bodies, which characterize the elongate-body morph, are often adaptations to open water environments, as has been supported by theoretical predictions for fish adapted to high sustained swimming (Webb, 1984; Ornelas-García et al., 2018). Also, the acute angular shape of the premaxilla in the elongate-body morph resembles the premaxillary shape found in other predatory characiforms, such as species of the genus *Oligosarcus* (Ribeiro & Menezes, 2015). Diversification related to trophic specialization has generally been recognized as a common pattern in characiforms (Sidlauskas, 2007, 2008; Ornelas-García et al., 2018).

The trophic niche of the deep-body morph inhabiting the lacustrine regions (lakes Managua and Nicaragua) was wider than that of the elongate-body morph. This pattern is congruent with results observed for similar morphs in other lacustrine populations of *Astyanax* (i.e., Lake Catemaco, Mexico) (Ornelas-García et al., 2018), suggesting trophic specialization in the elongate-body morph and retention of generalist habits in the deep-body one. Jepsen and Winemiller (2002) reported similar patterns for other fish species in Venezuela. In their analysis of stable isotope ratios, these authors showed that highly specialized piscivorous fishes, regardless of species, had less variable carbon and nitrogen profiles compared with other trophic groups. Thus, our finding of a narrower trophic niche for the lacustrine elongate-body morph is congruent with Bussing’s (1993) description of the species as having a primarily piscivorous feeding strategy. The wider trophic niche for the lacustrine deep-body morph suggests an omnivorous strategy, as found in other species of *Astyanax* (Mise, Fugi, Pagotto, & Goulart, 2013; Ornelas-García et al., 2018). In contrast to the lacustrine populations, in the Sarapiquí River, the elongate-body morph had a wider trophic niche compared with the deep-body morph, mainly due to the δ13C range. Despite our small sample size in some groups, the δ13C values could reflect the abundance of primary carbon sources within the food web, given that the range increases as different primary carbon sources are incorporated in the food web (Jackson et al., 2011). Differences in the direction of the niche amplitude observed between *Astyanax* morphs from the river and the lakes may be due to differences in primary carbon sources, and their dynamics, between the two environments. In lakes, pelagic and benthic primary carbon sources are known to contribute to higher trophic levels production (Vander Zanden & Vadeboncoeur, 2002); however, river food webs are considered more complex (Finlay & Kendall, 2007). River drainages contain both allochthonous carbon, derived primarily from terrestrial sources (plants and soils), and autochthonous carbon, derived from production within rivers (aquatic macrophytes, algae, and bacteria) (Finlay & Kendall, 2007). Further characterization of the food web in the Sarapiquí River and the Nicaraguan great lakes would likely clarify the relationship among primary carbon sources, the organisms that occupy several trophic levels, and the trophic niche width of morphs.

Likewise, the accumulation of N isotopes can be used to estimate trophic position as it increases with trophic level, with predators typically showing higher levels of accumulation compared with non-predators (Post, 2002; Jackson et al., 2011). We observed a positive correlation between N isotope accumulation and not only slender bodies but also acuter angles of the premaxilla, further support the idea that elongate-body morphs are predatory. We also compared the trophic level of sympatric morphs in the three regions relative to their detailed morphology. In our models, the factors related to the divergence between morphs (RW1 of body/premaxillary shape and morph type) were significant, showing evidence of divergence in trophic level. Interestingly, the differences in trophic level between sympatric morphs were robust and consistent, despite the differences in δ15N among regions, which could be explained by variation at the base of the food web from which organisms draw nitrogen (Post, 2002). Ornelas-García et al. (2018) also found a higher trophic level in the elongate-body morph compared with the deep-body morph in Lake Catemaco, México. Our results, together with those ofornelas-García et al. (2018), support the hypothesis that morphology could reflect the ecology of an organism, and could be used to predict its trophic habits, as has been proposed in other characids (Bonato, Burress, & Fialho, 2017).

### Genetic differentiation between morphs

We found slight genetic differentiation between sympatric morphs of *Astyanax* in the San Juan River basin, but with some contrasting patterns between morphs and regions. Our phylogenetic reconstruction of *Astyanax* did not recover monophyletic groups according to morph type or region, though the AMOVAs for both types of markers were congruent in showing differences among regions, supporting the possibility of some geographic genetic structuring. However, the AMOVA for the nuclear markers revealed significant differences between morphs when regions were considered as groups (i.e., Lake Managua, Lake Nicaragua, and Sarapiquí River), indicating some genetic divergence between morphs.

Furthermore, in the Fst pairwise comparisons, we found differences between sympatric morphs in the Sarapiquí River region, although comparisons between morphs in the lakes Managua and Nicaragua were not significant. A similar pattern of genetic differentiation has been described for the African cichlid fish *Astatotilapia burtoni*, in which the degree of differentiation between lake and stream adapted fish varied across regions in Lake Tanganyika (Theis, Ronco, Indermaur, Salzburger, & Egger, 2014). This pattern of varying genetic differentiation between morphs in different regions suggests that species might be at different stages of the speciation continuum (Theis et al., 2014). According to the clustering analyses, our samples most likely represent two (Bayesian clustering algorithm, Structure) or three (DAPC) populations, however, the clusters were not differentiated by morph type or region. Bayesian clustering analyses are known to have a low capability to detect distinct clusters when genetic differentiation levels are low (e.g., Latch, Dharmarajan, Glaubitz, & Rhodes, 2006). In cases of incipient speciation, genetic differentiation is typically weak (Nosil et al., 2009); therefore, our case may be reflecting a similar situation. Although Jombart et al. (2010) showed that DAPC generally performs better than Structure at characterizing population subdivision, and can even be used to unravel complex population structures, in our case, it did not detect any clustering associated with morphs or regions. Only the analyses based on the nuclear markers (AMOVA and Fst comparisons) provide evidence supporting slight genetic differentiation between morphs, and particular in the Sarapiquí River. Therefore, our study does not provide clear evidence of any gene flow limitations between morphs. Future studies should focus on determining if the significant genetic differentiation between morphs from Sarapiquí River is stable over time, particularly considering our study’s low sample size of elongate-body morphs from that region, where their presence is considered rare (Bussing 1998). Also, future studies should include more molecular markers (e.g., genomic data) to uncover potential differentiation patterns between morphs not detected by our data set.

Based on the weak genetic differentiation between morphs, we cannot rule out the alternate evolutionary scenario of stable polymorphism. Since the genetic basis of the ecomorphological divergence between morphs in the San Juan River basin is unknown, phenotypic plasticity (i.e., alternative phenotypes in response to environmental differences) remains as a plausible explanation for the adaptive divergence found, according to a stable polymorphism scenario (Smith & Skúlason, 1996). However, there is increasing evidence suggesting that phenotypic plasticity plays an important role in the ecological speciation process (Nonaka et al., 2015; Sabino-Pinto et al., 2019).

The evolution of barriers to gene flow (leading to reproductive isolation) and morphological divergence could be time-dependent, such that recently diverged populations could show weaker signs of genetic differentiation (under an ecological divergence framework) than older populations (Nosil et al., 2009; Nosil, 2012). The low level of genetic differentiation observed between sympatric morphs in the San Juan River system is higher than that recently reported in an equivalent system in Mexico (Lake Catemaco, Ornelas-García et al., 2020). In that system, no genetic differentiation was found between the elongate and deep-body morphs, and only a difference in migration rates between morphs was observed, tentatively suggesting a genetic differentiation cline across the Mexican and Central American lakes. The *Astyanax* found in Lake Catemaco may correspond to a younger lineage of the genus than that in the San Juan River basin (Ornelas-García et al., 2008), which could represent an older diverging lineage with discrete levels of genetic differentiation. Congruent with this genetic divergence, Garita - Alvarado et al. (2018) and Powers et al. (2020) found a higher level of morphological divergence between morphs in the San Juan River system than between those from Lake Catemaco. Additionally, besides time, Nosil et al. (2009) noted other factors that could influence the ecological speciation process including the nature of the divergent selection itself (e.g., strong vs. weak selection, or selection on a single trait vs. selection on a larger number of traits) and genetic factors (e.g., no physical linkage between genes under selection and those conferring reproductive isolation). These factors should be addressed in *Astyanax* from Lake Catemaco and the San Juan River basin to better understand the ecomorphological and genetic divergence between morphs.

### Isolation by distance

A pattern of isolation by distance was observed within the San Juan River basin. Geological evidence shows that Managua and Nicaragua lakes originally formed a single water body (the “Great Nicaraguan Lake”) that split into the present lakes in the late Pleistocene (Hayes, 1899; Villa, 1976). Today, the two lakes remain partially isolated, and only intermittently connect with each other through the Tipitapa River (see Fig 1), which runs partially subterranean, but experiences periodic surface water flow. This situation implies that contemporary exchange of fish between the two lakes is not extensive (Villa, 1976; Barluenga et al., 2010) and thus gene flow is limited by distance. Consistent with this are the results of the AMOVAs for both types of markers, which showed significant differences among regions. The complex (geo) hydrological history of this area could be reflected in the phylogenetic relationships recovered (the two lineages present in the San Juan River basin also include *Astyanax* from outside the basin), and in the observed pattern of isolation by distance. Further studies with additional samples could shed light on potential secondary contact or incomplete lineage sorting patterns in *Astyanax* in the San Juan River basin system.

In conclusion, we observed a strong correlation between body shape and premaxillary shape, highlighting the morphological divergence of a relevant trophic structure in sympatric morphs of *Astyanax* in the San Juan River system. We also found a strong correlation between this morphological divergence and divergence in trophic ecology (trophic niche and level) in these morphs, strongly supporting their adaptive divergence. Furthermore, the weak genetic differentiation observed between the morphs suggests they may be at a stage of incipient ecological speciation. Future studies including genomic data may shed light on the genetic basis of ecomorphological divergence in this model and its speciation process.

---

**Figure S1.**
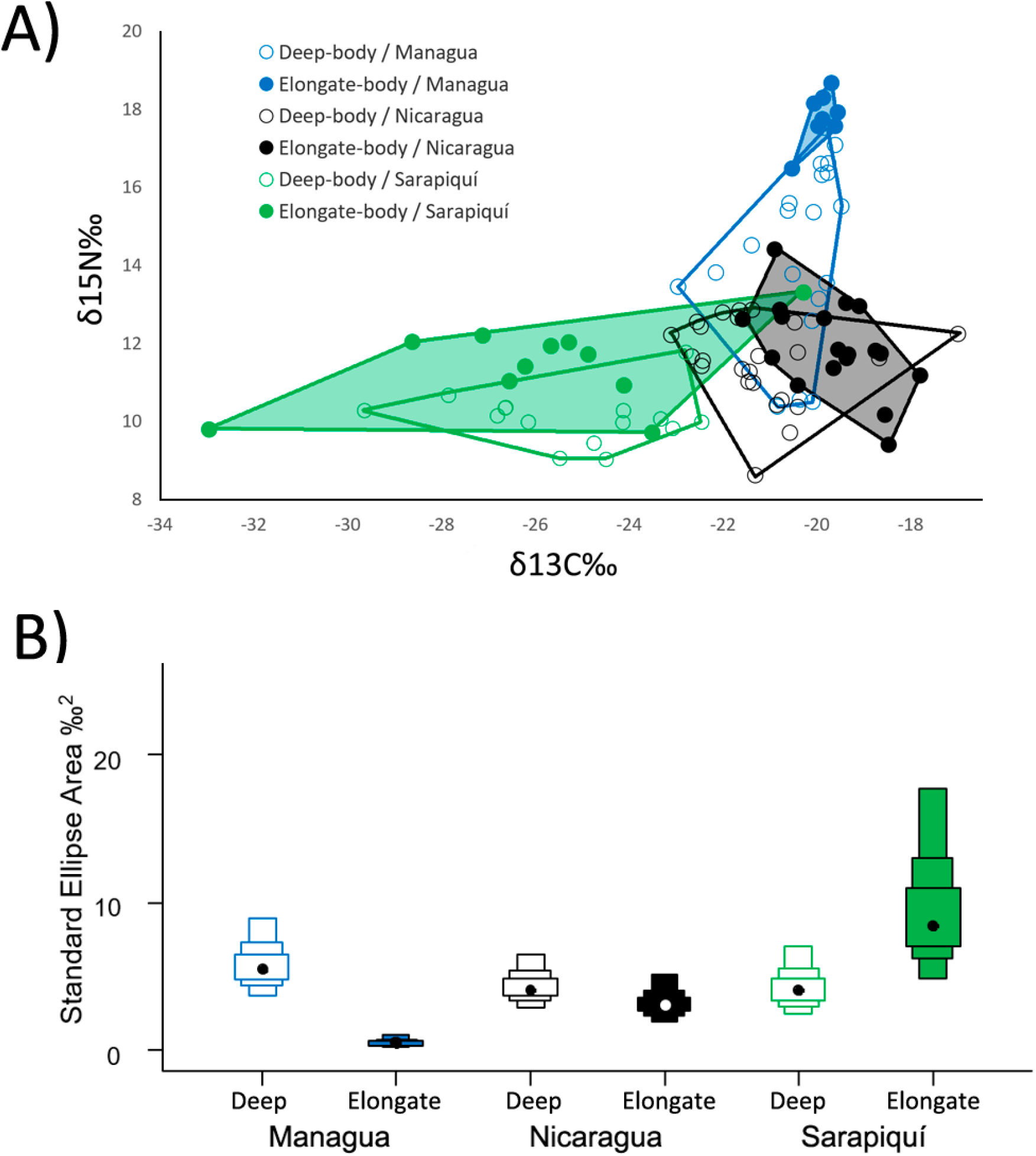
δ15N and δ13C plot (A), and Bayesian standard ellipse area by region and morph type (B). Empty boxes: deep-body morph, Solid boxes: elongate-body morph.

**Figure S2.**
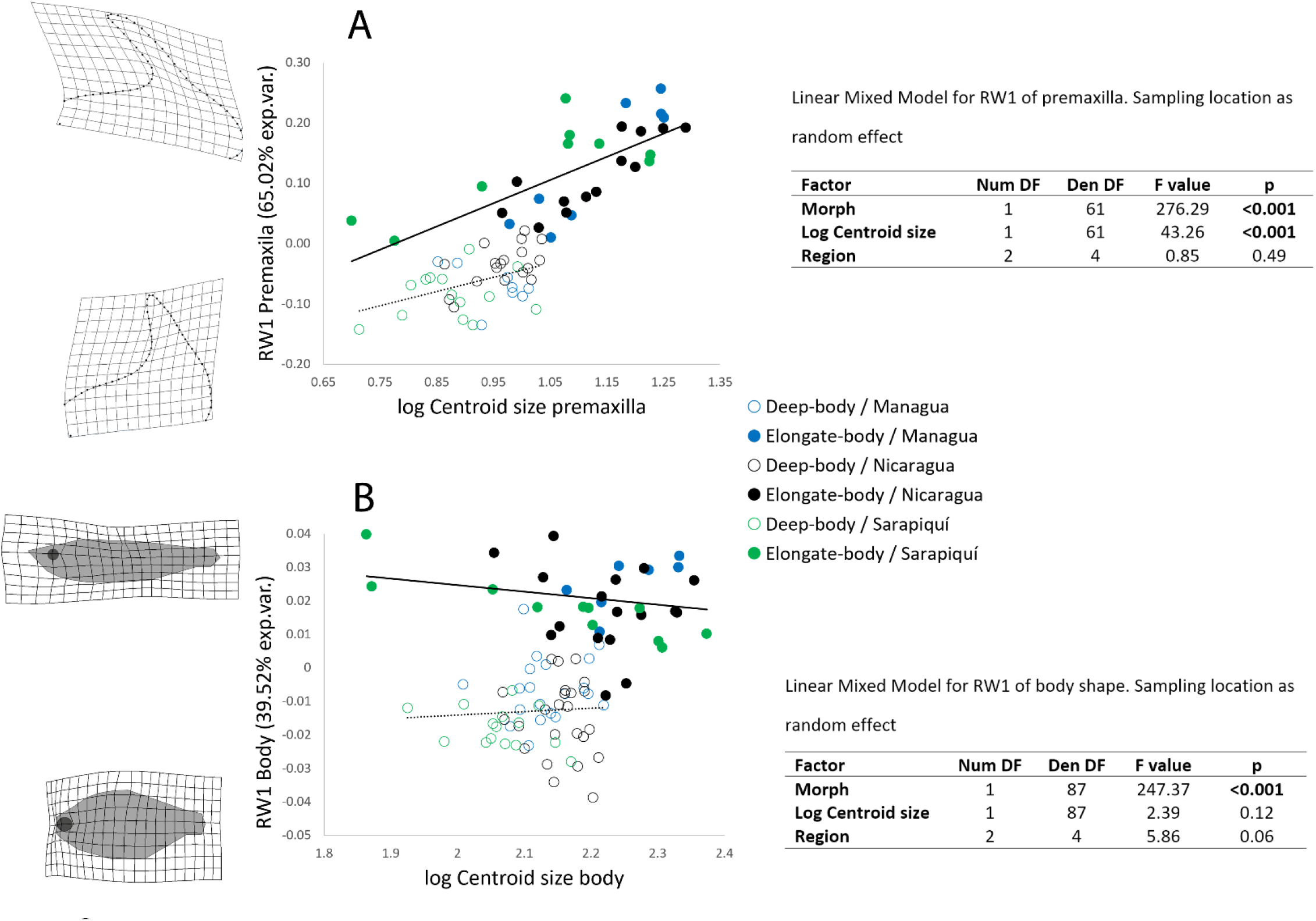
Variation in the shape of the premaxilla (A) and the body (B) associated with log centroid size. Results of the linear mixed models for each data set are shown on the right.

**Figure S3.**
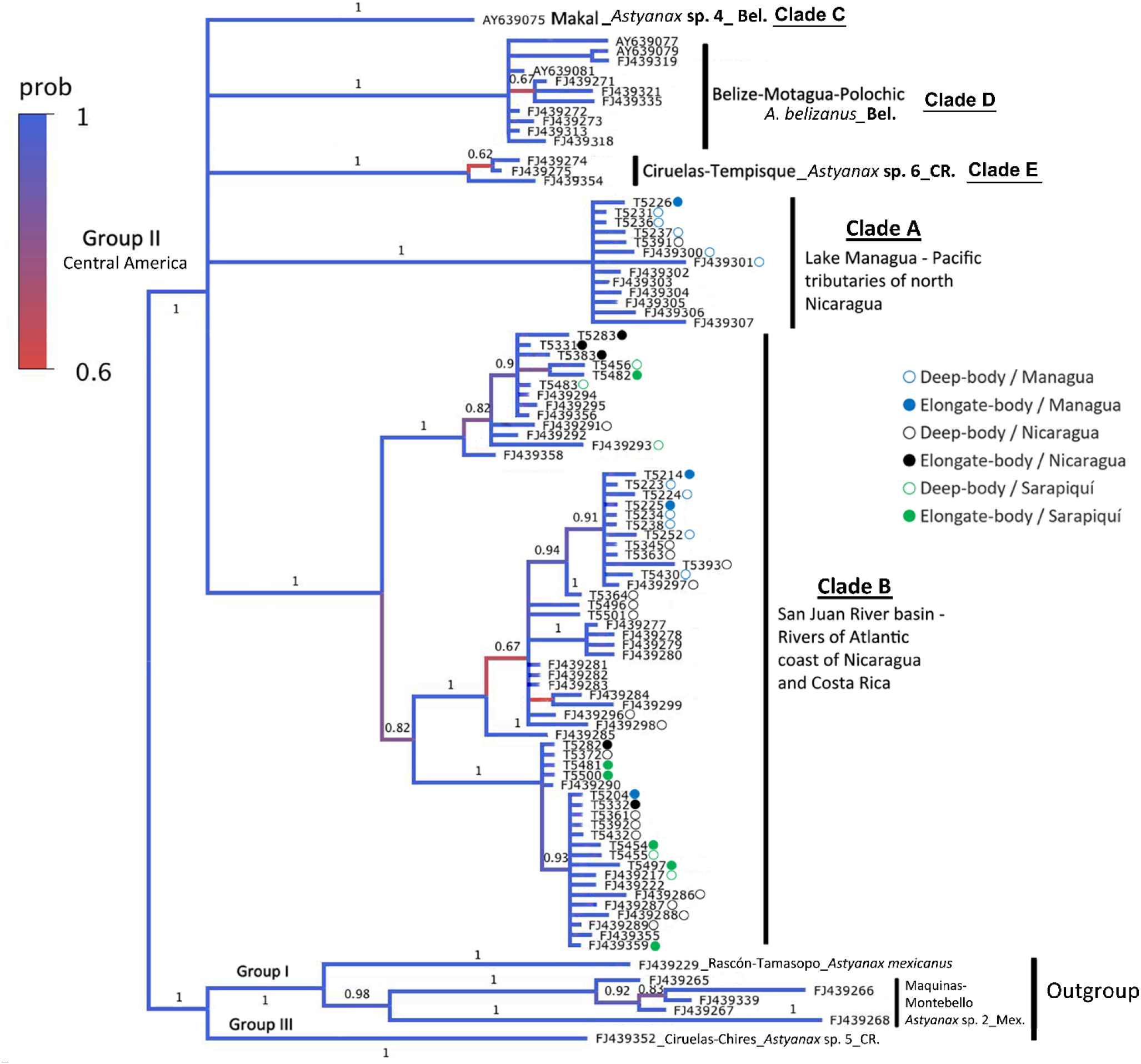
Bayesian phylogenetic tree (n=96) based on Cyt*b* for *Astyanax* in Central America. Sequences used include those of the Group II clade of *Astyanax* identified by Ornelas-García *et al*. (2008) and the 37 newly obtained for this study. Morph type is indicated for the samples from the three regions of the San Juan River basin. The posterior probabilities values are shown above branches (Bel = Belize, CR = Costa Rica, Mex = Mexico).

**Figure S4.**
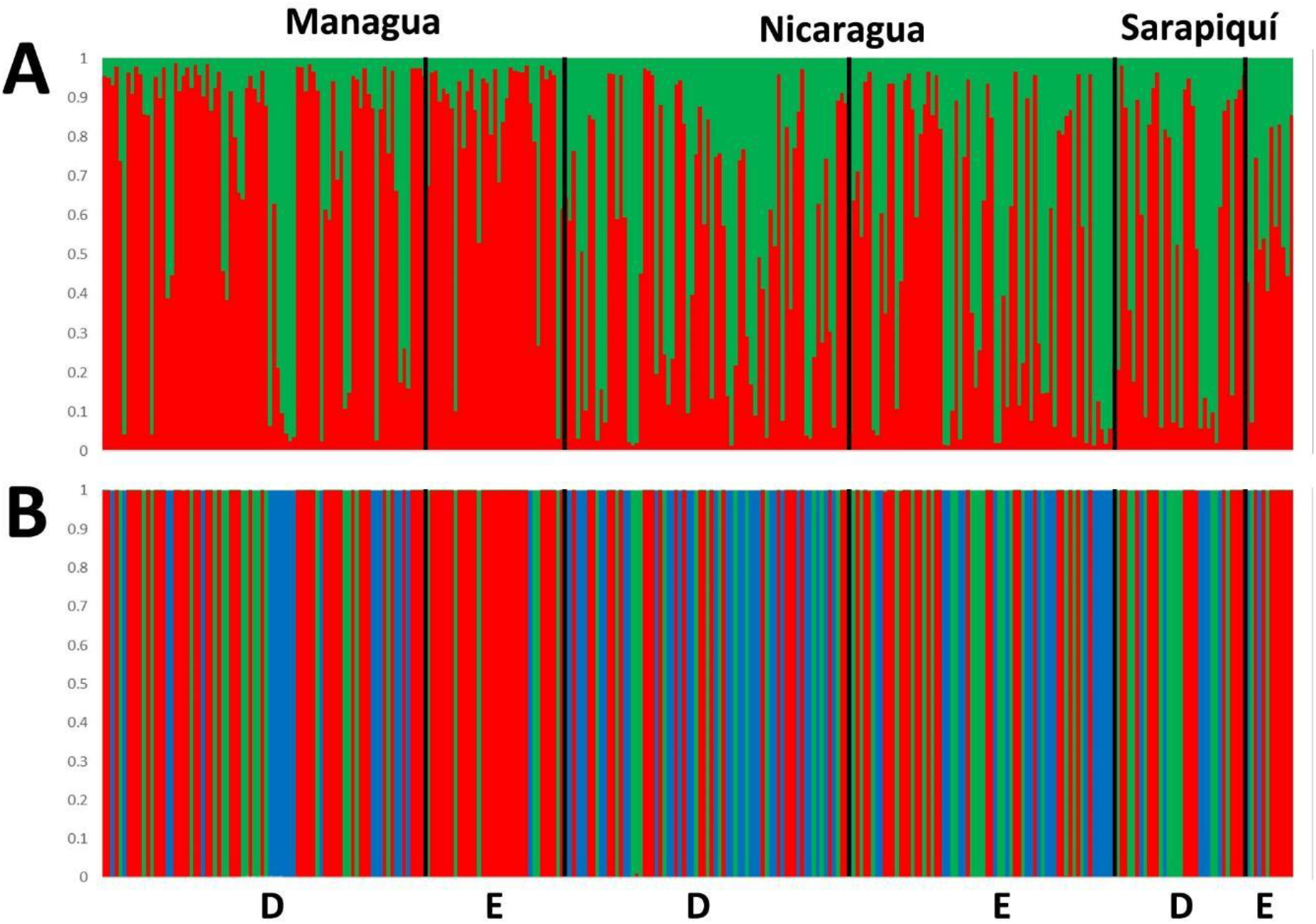
Clustering of individuals as determined by Structure (K = 2) and DAPC (K = 3). (D = deep-body morph, E = elongate-body morph).

**Figure S5.**
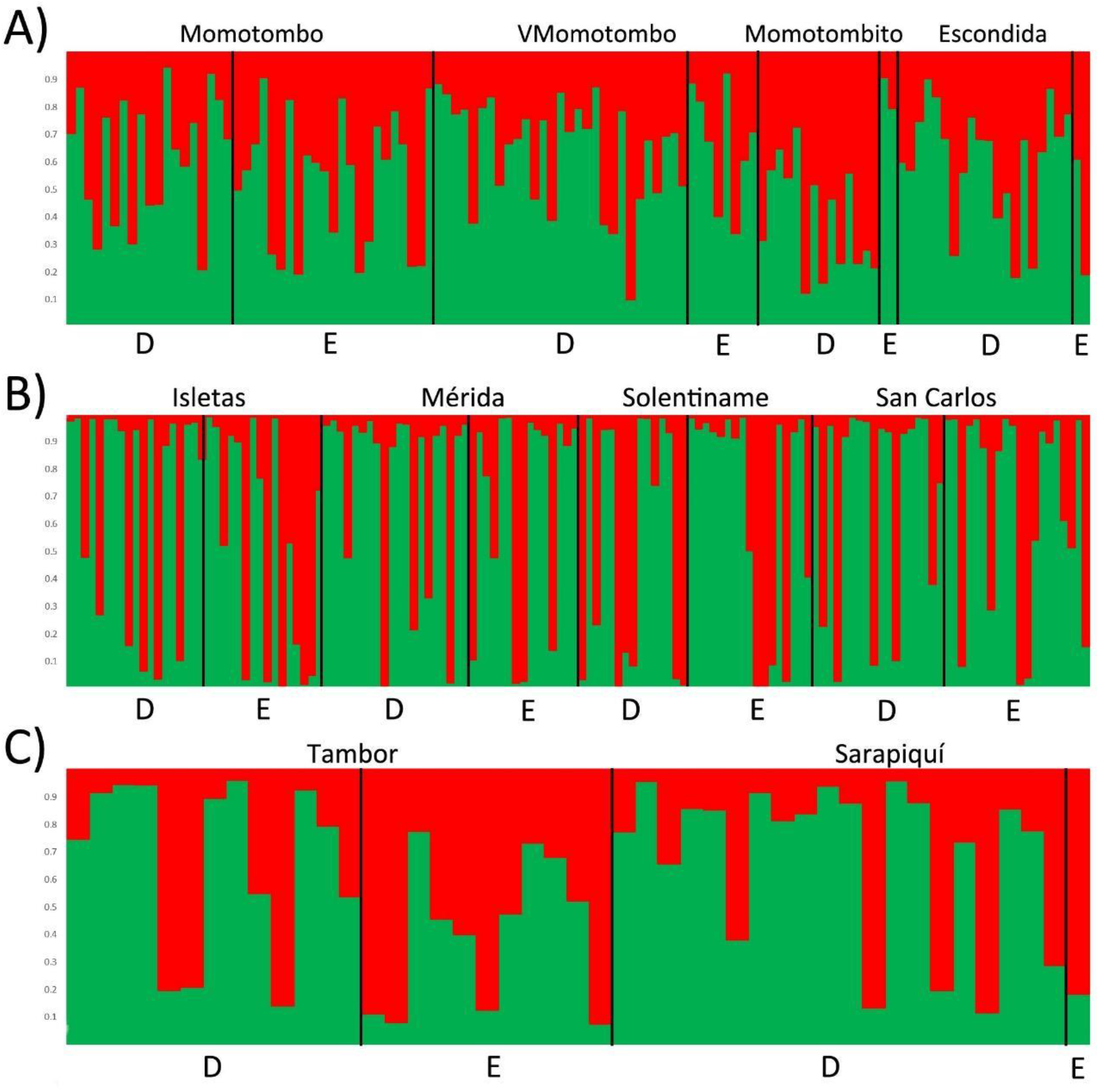
Clustering of individuals by region as determined by the Structure analysis showing K = 2 for A) Lake Managua, B) Lake Nicaragua, and C) Sarapiquí River. Sampling locations are indicated above each clustering. (D = deep-body morph, E = elongate-body morph).

**Table S1.** Layman (2007) metrics by region and morph. Total area: TA, mean distance to centroid: CD, mean nearest neighbor distance: NND, standard deviation of nearest neighbor distance: SDNND.

| Morph | $\delta^{15}\text{N}$ range | $\delta^{13}\text{C}$ range | TA | CD | NND | SDNND |
| --- | --- | --- | --- | --- | --- | --- |
| Deep-body Nicaragua | 4.25 | 6.11 | 13.15 | 1.41 | 0.47 | 0.49 |
| Elongate-body Nicaragua | 5.02 | 3.78 | 9.18 | 1.31 | 0.52 | 0.41 |
| Deep-body Managua | 7.15 | 3.48 | 13.38 | 2.07 | 0.46 | 0.27 |
| Elongate-body Managua | 2.20 | 0.97 | 0.78 | 0.53 | 0.40 | 0.34 |
| Deep-body Sarapiquí | 2.75 | 7.19 | 10.69 | 1.84 | 0.63 | 0.56 |
| Elongate-body Sarapiquí | 3.58 | 12.67 | 23.73 | 2.34 | 1.53 | 1.62 |

